# Structure of SARS-CoV-2 spike in complex with its co-receptor the neuronal cell adhesion protein contactin 1

**DOI:** 10.64898/2026.03.28.714969

**Authors:** Sabrina T. Krepel, Daniel L. Hurdiss, Berend J. Bosch, Joost Snijder, Bert J.C. Janssen

**Affiliations:** Biomolecular Mass Spectrometry and Proteomics, Bijvoet Center for Biomolecular Research and Utrecht Institute of Pharmaceutical Sciences, Utrecht University, Utrecht, The Netherlands; Structural Biochemistry, Bijvoet Center for Biomolecular Research, Department of Chemistry, Faculty of Science, Utrecht University, Utrecht, The Netherlands; Virology section, Infectious Diseases and Immunology Division, Department of Biomolecular Health Sciences, Faculty of Veterinary Medicine, Utrecht University, The Netherlands

## Abstract

The emergence of SARS-CoV-2 has caused millions of deaths and excess morbidity in the worldwide population. In addition to its respiratory symptoms, SARS-CoV-2 has become known for its neurotropism and long-term neurological sequelae, with a post-acute infection syndrome commonly referred to as long-COVID. Next to the host receptor angiotensin-converting enzyme 2 (ACE2) additional interactions of the SARS-CoV-2 spike (S) protein have been described for neuronal co-receptors specific to the nervous system including cell adhesion protein contactin 1 (CNTN1). Details of the spike-CNTN1 interaction have remained elusive. Here, we quantified the spike-CNTN1 interaction by surface plasmon resonance and resolved the structure of the complex by single particle cryo-electron microscopy (cryo-EM). Spike and CNTN1 interact with nanomolar affinity, driven by an avidity effect and mediated by the horseshoe moiety of CNTN1. The cryo-EM structure reveals that the CNTN1 Ig1-4 horseshoe is wedged in between two receptor binding domains (RBDs) and interacts, through Ig3, with a unique receptor interface at the base of the RBD in the up-conformation. This receptor interface is not previously described for other spike receptors but overlaps with the epitopes of several neutralizing monoclonal antibodies. Comparison of our data with available spike structures suggests one spike trimer can bind three CNTN1 molecules, or alternatively, different co-receptors such as ACE2 and CNTN1, simultaneously. These findings shed new light on the molecular determinants of SARS-CoV-2 neurotropism.

## Introduction

The emergence of SARS-CoV-2 in recent years has caused excess mortality and morbidity in the global human population^1^. While the virus is known primarily for its respiratory symptoms, the 2019 pandemic and subsequent seasonal epidemics have also brought to light the neurotropism of SARS-CoV-2 and its long-term neurological sequelae, with a post-acute infection syndrome known as long-COVID^2–8^. Because of the prevalence of neurological symptoms in severe COVID-19 and long-COVID, neuroinvasion and neurovirulence has been extensively studied for SARS-CoV-2^7,9–12^. Infection has been shown in neurons, astrocytes and vascular endothelial cells in the central nervous system^13–17^, which may result from infection via cranial nerves such as the trigeminal ganglion or olfactory nerve^18–21^, or by infection via the hematogenous route through a weakened blood-brain barrier (BBB)^22–24^. In addition, factors beyond neuroinvasion might be responsible for the neurological complications of SARS-CoV-2, with the relative contributions of neurotropism or systemic inflammatory responses subject of ongoing research^21–23,25^.

The host range and tissue tropism of SARS-CoV-2 is largely determined by specific interaction of the viral envelope membrane bound spike protein (S) with host receptors. Spike is a trimeric class I fusion protein with each protomer consisting of an S1 and S2 subunit, which are separated by a furin cleavage site^26,27^. The N-terminal S1 subunit includes the N-terminal domain (NTD) and the receptor-binding domain (RBD), of which the latter exhibits dynamic changes between an exposed upward or protected downward conformation, allowing for receptor binding or immune shielding, respectively. Upon receptor binding, the S1 domain dissociates from the S2 domain, yielding the metastable prefusion state of S2 which transitions to its stable postfusion state to drive fusion of the viral envelope and host cell membrane^27–31^.

Various host factors have been identified that can interact with SARS-CoV-2 in the nervous system. The primary host receptor of SARS-CoV-2, angiotensin-converting enzyme 2 (ACE2), interacts with the viral envelope via the fusion spike (S) protein, which triggers membrane fusion^32,33^. Additional spike interactors have been identified that facilitate infection in both ACE2-dependent and -independent ways, such as neuropilin-1 (NRP1)^17,34–36^, TMEM106B^37^, tyrosine-protein kinase receptor UFO (AXL)^38^, dipeptidyl-peptidase 4 (DPP4 or CD26)^16^, basigin (CD147)^16^, Siglec-9^39,40^, and contactin 1 (CNTN1), for which many molecular details of the interaction with spike remain to be explored^41,42^.

CNTN1 is a neuronal cell adhesion protein highly expressed in the CNS^41,43^ and involved in several developmental processes including neurological cell differentiation, myelination, synapse formation and microglia activation through interleukin expression. It is mainly expressed by oligodendrocyte precursor cells (OPC), astrocytes and neurons, where it forms adhesion complexes with neurofascin and caspr1 to regulate the paranodal connection^44,45^. CNTN1 consists of 6 N-terminal immunoglobulin(Ig)-like domains and 4 fibronectin type III (FNIII) domains, which are C-terminally glycophosphatidylinositol (GPI) anchored into the membrane^43,46^. Several N-glycosylation sites, primarily located in the Ig1-6 domains, decorate CNTN1 with glycans involved in stabilizing complex formation with neurofascin. The N-terminal Ig1-4 domains form a horseshoe arrangement characteristic for a subset of members of the IgCAM family. In an interaction screen, CNTN1 was identified as a novel RBD-binding co-receptor, specific for SARS-CoV-2 with no apparent SARS-CoV-1 RBD binding^41,42^. Expression of CNTN1 with ACE2 on HEK293T cells enhanced infection of SARS-CoV-2 pseudotyped particles, with an even greater effect when TMPRSS2 was also present^41^. The ACE2-dependent enhancement of infection by CNTN1 expression surpassed the effect of NRP1, which further emphasizes the potential role of CNTN1 to be an important regulator of SARS-CoV-2 neurotropism.

Here, we studied the molecular details of the SARS-CoV-2 spike-CNTN1 interaction. We quantified the binding strength of trimeric spike with CNTN1 by surface plasmon resonance and pinpoint the interaction to the Ig-domains of CNTN1. We provide further details of the interaction by single particle cryo-EM analysis of a complex between the spike trimer and the full extracellular segment of CNTN1. The structure reveals that the spike RBD in the up-conformation interacts directly with the Ig3 domain in the horseshoe of CNTN1. We discuss important differences and similarities with other SARS-CoV-2 host receptor interactions and compare the spike-CNTN1 interface within SARS-CoV-2 variants and SARS-CoV-1. Our findings provide a structural basis to understand the molecular determinants of SARS-CoV-2 neurotropism and may help to better understand disease mechanisms responsible for the virus’ long-term neurological sequelae.

## Results

### SARS-CoV-2 spike binds to CNTN1 with nanomolar affinity through its Ig-domains

SARS-CoV-2 prefusion stabilized spike (6P) and CNTN1 interact with a K_D_ of 67 ± 12 nM in an SPR equilibrium binding experiment (Fig. 1A-D). To mimic the native cell-surface attachment topology of CNTN1 we immobilized CNTN1 via its C-terminus to the sensor surface and used the spike trimer as analyte. The equilibrium data was initially fitted to a 1:1 Langmuir model (Fig. 1C,D). At higher concentrations of spike (>250 nM), a second, lower affinity interaction is apparent. When including the high concentration equilibrium points in our model (Fig. E,F), these data fit well to a two-state specific binding model that indicates a high affinity interaction (52 nM, K_D1_), matching the analysis shown in Figure 1C-D, and a lower affinity interaction (12 µM, K_D2_) (Fig. 1E,F, Supl. Fig S1). Possibly, these data reflect a high affinity multivalent interaction (K_D1_) in which one spike trimer interacts with two or three CNTN1 molecules on the sensor surface and a monovalent interaction (K_D2_) in which a spike trimer interacts with a single CNTN1 molecule. In addition, we found that interaction of spike with CNTN1 is dependent on the presence of its 6 N-terminal Ig-domains (Fig. 1E). Binding is abolished for a CNTN1 construct lacking the first six Ig domains (CNTN1^FNIII^, Fig. 1E).

**Figure 1.**
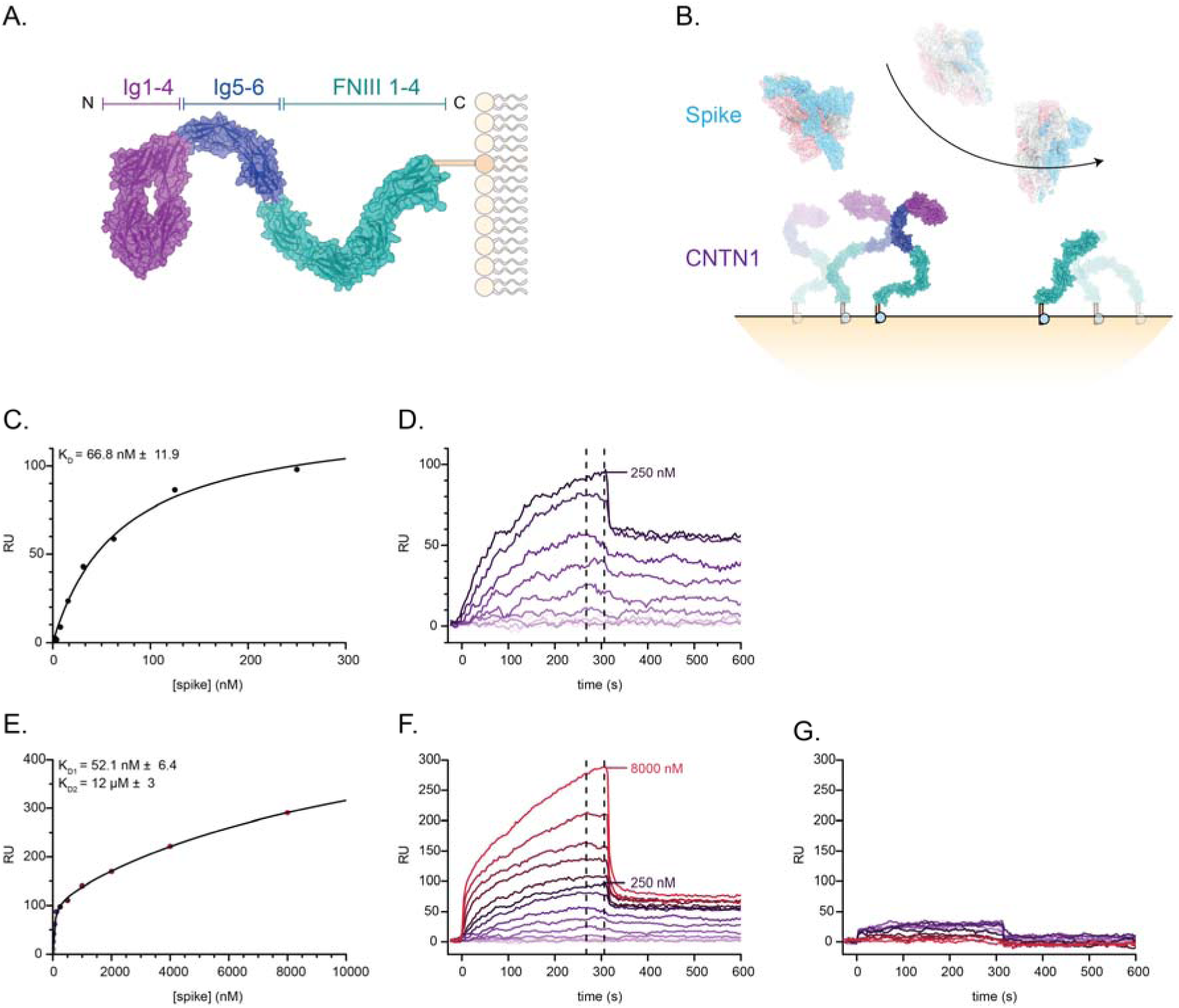
SARS-CoV-2 spike binds CNTN1 with a K_D_ of 67 nM. A. Domain organisation of CNTN1. B. Experimental set-up of SPR measurements where trimeric prefusion stabilized Wuhan spike is used as analyte and CNTN1 constructs used as ligands. C. 1:1 Langmuir model fitting of equilibrium data based on sensogram data. D. Binding of spike to CNTN1 at concentrations ≤250 nM. E. Two-state binding model fitting to equilibrium data based on sensogram data of F. F. Binding of spike to CNTN1 at concentrations ≤8 μM. Note that at the highest concentrations of spike equilibrium is not reached. G. SPR sensogram data of spike with CNTN1^FN1–4^, no interaction is apparent up to a concentration of 8 µM spike.

### SARS-CoV-2 spike interacts with CNTN1 Ig3 via its RBD in the upward conformation

We determined the cryo-EM structure of a spike-CNTN1^fe^ complex, revealing one SARS-CoV-2 spike trimer interacting with one CNTN1 molecule. In the complex, the tip of the CNTN1 Ig1-4 horseshoe is wedged in between two RBD domains of spike, one of which is in the upward conformation (RBD^up^) and the other in the downward conformation (RBD^down^) (Fig. 2A-C, Suppl. Fig. S2). While the nominal FSC resolution of the map was determined at 3.5 Å (Table 1, Fig. S2), the RBD^up^-CNTN1^Ig1–4^ part was less well-resolved, likely due to substantial flexibility relative to the main body of spike. Focussed refinement using a mask encompassing only the RBD^up^ and Ig1-4 of CNTN1 improved its local resolution to an estimated 7.2 Å. Although this resolution prevents us from performing a detailed analysis of the interface at sidechain level, the placement of the CNTN1 Ig1-4 domains and RBD^up^ could be determined confidently and was aided by the availability of a higher resolution structure of CNTN1^Ig1–4^, ^46^, guided by low-resolution EM densities consistent with two N-linked glycans in CNTN1 (Fig. 2B). The flexible FNIII tail and Ig domains 5 and 6 of CNTN1 could not be resolved, indicating that this part is not involved in interactions with spike, in line with our SPR analysis (Fig. 1E).

**Figure 2.**
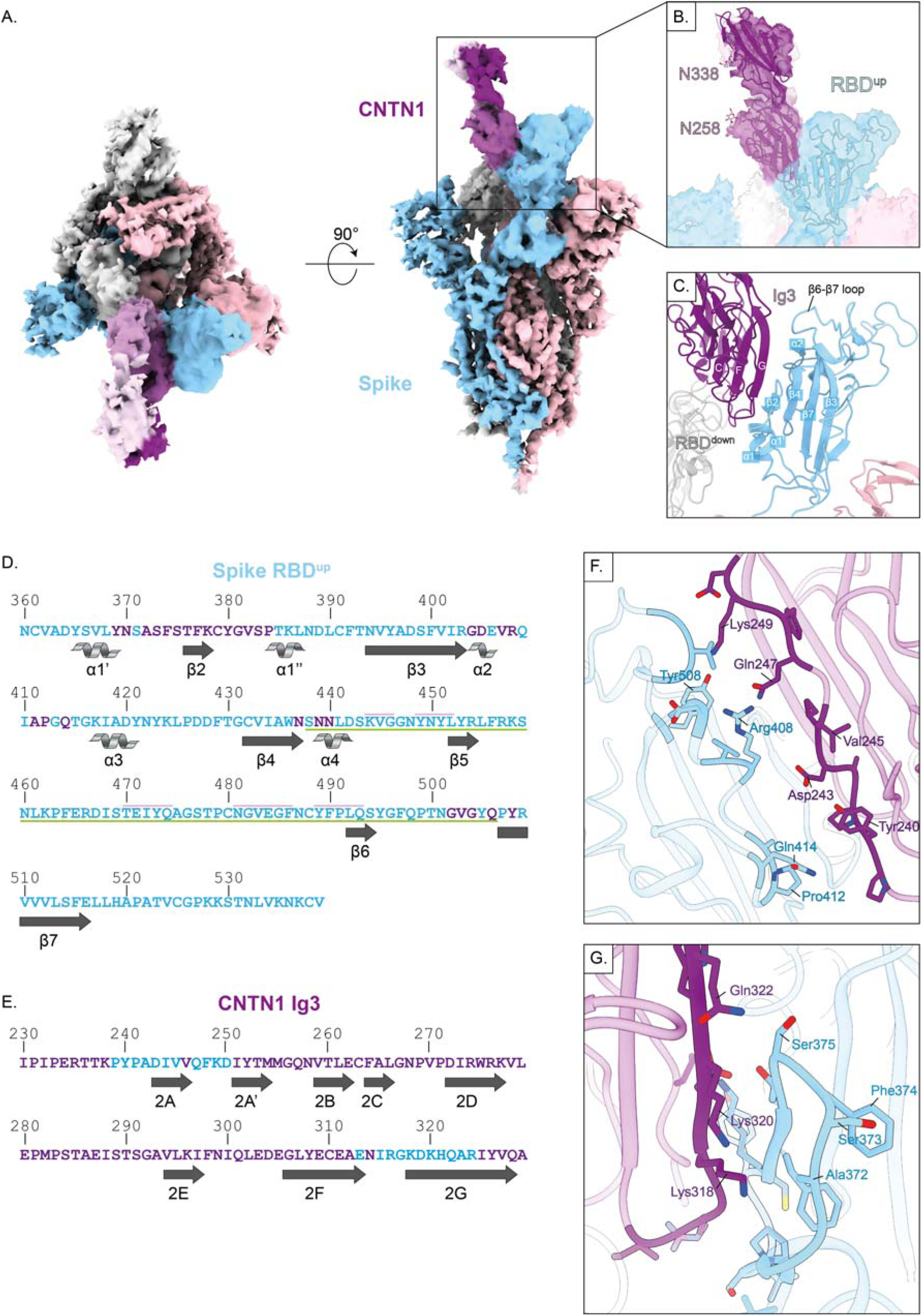
SARS-CoV-2 spike in complex with CNTN1. A. Cryo-EM structure of trimeric spike in complex with one CNTN1 molecule. spike protomers are coloured in light blue, pink and light grey, CNTN1 domains 1-4 are coloured in shades of magenta. B. Orientation of CNTN1 in the Cryo-EM map with glycan densities as guides. Glycan structures are obtained from the available crystal structure of CNTN1 (PDB ID: 7OL2^46^). C. Details of beta sheet orientation of CNTN1 Ig3 with regards to the RBD^up^. D. Secondary structure organization and contacts of RBD^up^ with CNTN1 (magenta), ACE2 (underlined) and TMEM106B^78^ (pink lines above). Secondary structure naming and ACE2 contacts are based on Lan et al^78^. E. Contacts and beta sheets of CNTN1 Ig3 with RBD^up^ (light blue). F. Residues of RBD^up^ β7 with CNTN1 beta sheet 2A. G. Residues of RBD^up^ β2 with CNTN1 beta sheet 2G.

The main intermolecular interface is formed between spike RBD^up^ and CNTN1^Ig3^. The RBD^up^-Ig3 interface buries a total surface area of approximately 1.500 Å^2^. RBD^up^ β-strand β2 and helixes α2 and α5 interact with the bottom half of Ig3 β-strands A and G that form the main part of the interface on the CNTN1 side (Fig. 2C-G). Simultaneously, one of the two RBD^down^ domains interfaces with parts of CNTN1^Ig2–3^. The buried surface area of the RBD^down^ – Ig2-3 interface is approximately 1.400 Å^2^ and involves loop EF and β-strand G of CNTN1 Ig2, and loop BC and FG of CNTN1 Ig3. Three-dimensional classification of the complex particles in two classes and subsequent refinement of the two sets shows that the RBD^up^ – CNTN1^Ig1–4^ combination moves in concert, relative to RBD^down^ (Suppl. Fig. S3). Consequently, in one of the two classes the interface between RBD^down^ and CNTN1^Ig1–4^ is reduced in buried surface area, while the RBD^up^ – CNTN1^Ig1–4^ interface is identical in both classes. This suggests that the RBD^up^ – CNTN1^Ig1–4^ interface plays the predominant role in the interaction of spike with CNTN1.

### The spike-CNTN1 interface is unique amongst known receptors and allows for multivalent interactions

Two different spike-receptor complexes for SARS-CoV-2 have previously been resolved by detailed structural analyses. Comparing the ACE2^47^ and TMEM106b^37^ receptor binding sites with the CNTN1 interface on spike shows that the CNTN1 binding site is not used by the two other receptors (Fig. 3). ACE2 binds to the RBD^up^ via the receptor binding motif (RBM) between the β4 and β7 strands including the short α4 and α5 helices and β5/β6 strands (Fig. 2D, green lines). This site is largely different to the CNTN1 binding site that consists predominantly of the β2 strand and the α2 and α5 helices. Only few of the contact residues on RBD^up^ overlap between CNTN1 and ACE2. TMEM106b interacts in a manner more reminiscent of ACE2 where the β6 strand defines most of the interacting surface to the posterior of RBD^up^ with respect to the ACE2 binding site. No interfacing residues on RBD^up^ overlap between CNTN1 and TMEM106b (Fig. 2D, pink lines).

**Figure 3.**
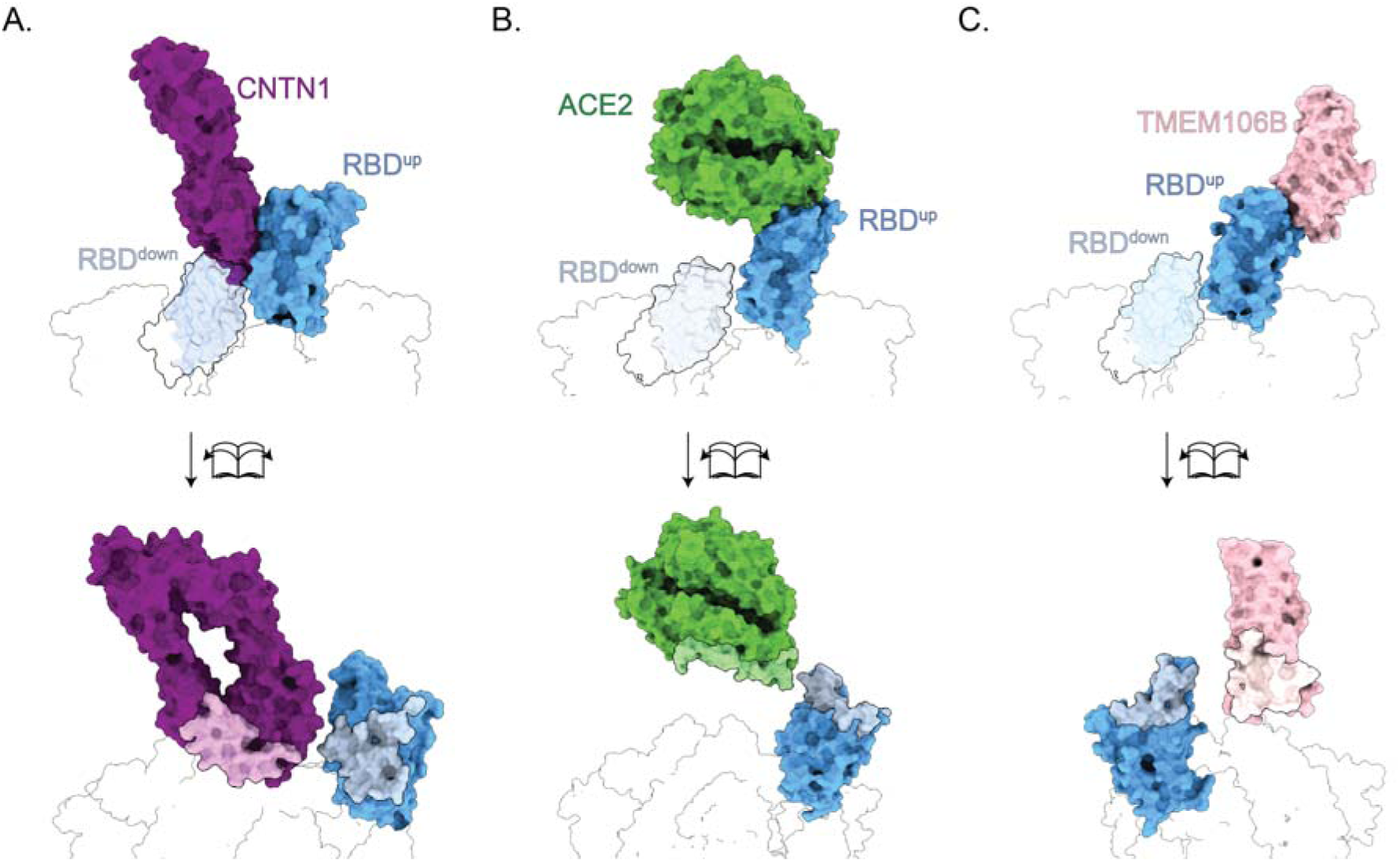
Binding interface of spike co-receptors to the RBD^up^ depicted in closed and open book representations. Interfaces were calculated with PDBePISA and depicted in lighter respective colours. A. Representation of the obtained CNTN1:spike model. B. Representation of ACE2 bound to the RBD^up^ (PDB ID: 6VW1^47^). C. Representation of TMEM106B bound to the RBD^up^. A structure of spike (PDB ID: 7NTA^79^) and TMEM106B (PDB ID: 8B7D^78^) were fitted in the density map obtained from the EMDB (EMDB ID: 17169^78^) in ChimeraX^69–71^.

Spike-CNTN1 complex formation does not appear to induce conformational changes in either of the proteins. The relative position of the four Ig domains in the CNTN1 horseshoe are the same compared to unliganded CNTN1^Ig1–6^,^46^. In addition, we observe the same upward conformation and position of the RBD^up^ in a subset of spike particles that are not in complex with CNTN1. Focussed 3D classification on all spike particles with one RBD^up^ (i.e. including those without CNTN1 bound) yielded volumes for six distinct conformations of RBD^up^ relative to the rest of the spike trimer, revealing a hinge-like motion for RBD^up^ (Suppl. Fig. S4). This flexibility of RBD^up^ has been observed before^30^. In two of the six classes, the RBD^up^ position is such that CNTN1 binding would be sterically hindered by the proximal RBD^down^, suggesting that the flexion in RBD^up^ is required for CNTN1 complex formation.

Multivalent interaction of CNTN1 binding to multiple RBD^up^ requires opening of multiple RBD domains. In our data, we observe approximately half of total spike particles with three RBD^down^ and half with one RBD^up^. Classes with more than one RBD^up^ were not identified. Notably, straightforward modelling by RBD-based superposing of the RBD-CNTN1 segment onto a spike structure where all three RBD’s are in the up conformation (i.e. PDB ID: 7R40^48^) shows no steric hindrance for binding of three CNTN1 molecules simultaneously (Suppl. Fig. S5). The spike binding site on CNTN1, at the tip of the horseshoe, is located furthest from the CNTN1 cell-surface binding site at the C-terminus of the last FNIII domain, providing ample room for multivalent interaction in a physiological setting. In addition, flexibility of CNTN1^Ig1–4^ with respect to Ig5-6 and the FNIII domains, apparent in our data and previously reported^46^, further extends the possibility of multiple CNTN1 molecules binding to one spike trimer. Thus, it is possible that a spike trimer can interact with two or even three cell-surface attached CNTN1 molecules, supporting the avidity effect we observe in our interaction data (Fig. 1).

The position of CNTN1 in relation to the RBD^down^ in our complex structure suggests that CNTN1 sterically hinders this domain to adopt the RBD^up^ position, thereby decreasing the likelihood of all RBD domains sequentially opening. On the other hand, the RBD of the third protomer, in the downward position, is not sterically hindered to open. CNTN1 binding to one RBD^up^ likely forces that RBD to stay in the upward position allowing the unhindered RBD^down^ to adopt the upward position by simple stochastics, i.e. the likelihood for the spike-CNTN1 complex to have two RBD^up^ is increased, although the frequency may remain low overall. If the second RBD^up^ also binds a CNTN1 molecule, sufficient room is created for the last RBD^down^ to transition to the upward conformation enabling three CNTN1 molecules to bind one spike trimer. Scenario’s where a spike trimer interacts with two different receptors seem also feasible from a structural perspective. Simultaneous CNTN1 and ACE2 interaction through different protomers within the same trimer should be possible (Suppl. Fig. S5), while both cannot bind the same spike RBD^up^ in the same protomer due to steric hindrance. Simultaneous interaction of CNTN1 and TMEM106b to the same RBD^up^ does not seem to lead to steric hindrance, although it is not clear if this can occur in a physiological setting.

### The spike-CNTN1 interface in SARS-CoV-1

Since it was previously established that SARS-CoV-1 spike does not interact with CNTN1^49^, we examined its differences with the CNTN1 binding site of SARS-CoV-2 (Fig. 4A). Most differences at residue level are at the ACE2 interaction site. Nonetheless, five differences are located within the CNTN1 interface on RBD^up^. Comparing SARS-CoV-1 with SARS-CoV-2 these are T372A, F373S, A384P, R439N and I503V, of which T372A and F373S cause a local reduction in hydrophobicity. The low resolution of our structure limits a more detailed analysis of the consequences of each mutation on CNTN1 interaction, and the question remains how much each substitution contributes to prevent interaction between SARS-CoV-1 and CNTN1. One other notable difference is the absence of an N-linked glycan at N370 in SARS-CoV-2 due to mutation of the NST motif in SARS-CoV-1 to NSA in SARS-CoV-2. A bulky N-linked glycan at this site would be placed near to the binding interface between CNTN1 and spike. However, superposition of SARS-CoV-1 RBD structures onto the RBD^up^ in the spike-CNTN1 complex show that the SARS-CoV-1 RBD glycan is oriented away from the CNTN1 binding site^50,51^ and is therefore unlikely to sterically hinder interaction.

**Figure 4.**
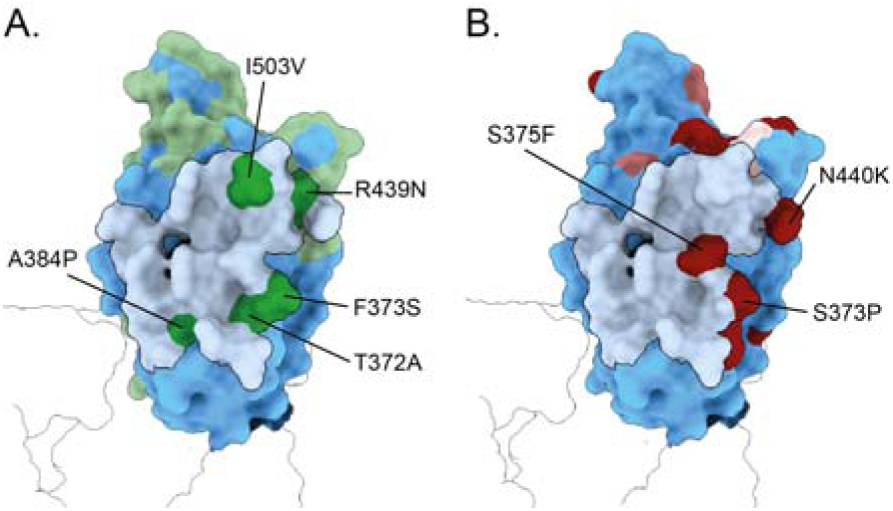
Comparison of SARS-CoV-2 Wuhan spike RBD with SARS-CoV-1 spike RBD and more recent variants. A. Representation of the RBD in the up conformation with outlined CNTN1 binding area (light blue) and highlighted differences with SARS-CoV-1 in green. Residue differences within the binding area are indicated. B. Representation of the RBD in the up conformation with variant mutations and binding area of CNTN1 highlighted. Lighter red colours indicate mutations that emerged earlier. Light red = alpha (B1.1.7), medium red = beta (B1.35) and gamma (P.1), dark red = omicron (BA.1). Mutations within the binding interface with CNTN1 are indicated.

### The CNTN1 binding site on RBD overlaps with class 4 neutralizing antibody epitopes

Numerous neutralizing monoclonal antibodies that target SARS-CoV-2 RBD have been reported and structurally characterized, commonly grouped into four main classes based on overlapping epitopes^52^. Among these, class 4 monoclonal antibodies have the biggest overlap with the CNTN1 binding site on RBD (Fig. 5). Notable examples include monoclonal antibodies CR3022^53^ and S304^54^, as well as nanobody VHH-72^55^, whose epitopes have substantial overlap with the CNTN1 footprint to produce severe clashes when simultaneously docked to the same RBD.

**Figure 5.**
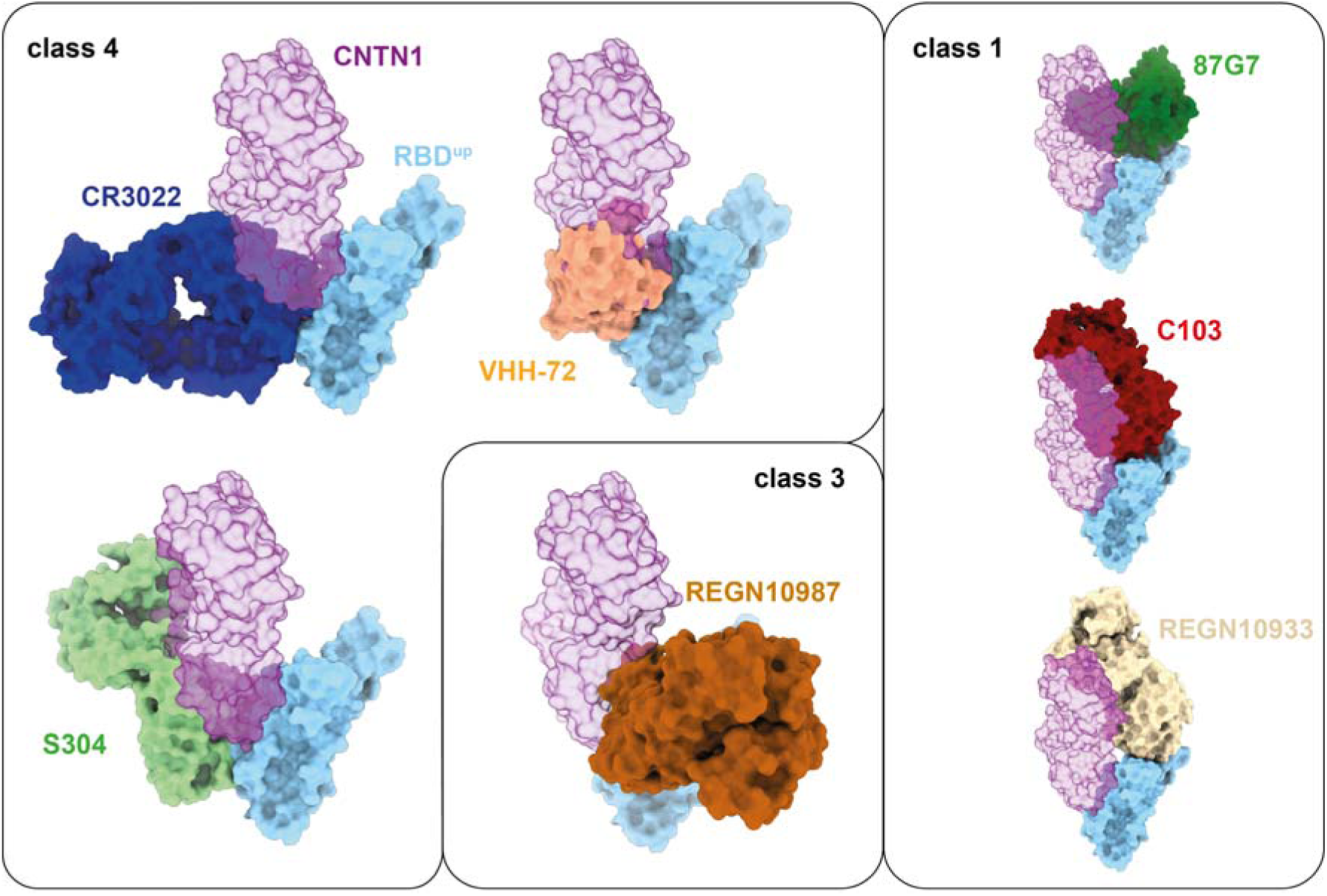
CNTN1 clashing antibody and nanobody classifications of SARS-CoV-2 spike. Epitopes of class 4 antibodies and nanobodies overlap with the CNTN1 binding site on RBD. For class 1 and 3 antibodies the binding sites are not overlapping but steric clashes are apparent between the antibodies and CNTN1.

Interestingly, while CR3022 binding destroys the native quaternary structure of spike^56^, we see no evidence of this in our spike-CNTN1 reconstructions, likely due the different angles of approach between CNTN1 and CR3022. Whereas CR3022 binding clashes with the neighbouring RBD protomer to disassemble the S1 subunits,

CNTN1 rather neatly wedges exactly within the available space between the binding RBD^up^ and neighbouring RBD^down^ subunits. While the other classes of neutralizing antibodies against SARS-CoV-2 RBD do not have any notable overlap with the CNTN1 footprint, members of the class 1 (87G7^48^, C102^52^, REGN10933^57^) or class 3 (REGN10987^57^) antibodies may still result in substantial clashes with CNTN1 due to the extended structures of their respective Fab and horseshoe domains.

## Discussion

Infection of SARS-CoV-2 can have detrimental consequences to the central nervous system, leading to life-altering symptoms such as headaches, dizziness and exhaustion, but also encephalitis and seizures^58^. Here, we confirm the interaction of CNS-specific co-receptor CNTN1, found only to interact with SARS-CoV-2 spike as opposed to SARS-CoV-1 spike. We quantified the interaction affinity by SPR to be 67 nM, which is enhanced by an avidity effect likely reflecting the ability of one spike trimer to bind to multiple CNTN1 molecules.

In a physiological context, avidity-driven affinity enhancement probably increases the ability of SARS-CoV-2 to bind to CNTN1 and utilize it as a co-receptor. Our cryo-EM structure of CNTN1-bound spike reveals an important binding site at the base of the RBD^up^, which is only available in its upward conformation. When all three RBD’s are in the downward conformation the three CNTN1 binding sites are largely hidden and, supposedly, shielded from the immune system. In contrast to ACE2 and TMEM106B, which bind at the tip of the RBD^up^ (Fig. 3B,C), the placement of CNTN1 at the base of RBD^up^ poses steric hindrance for opening of the adjacent RBD^down^ (Figure 3A), while the third RBD (i.e. the second in the down conformation) is unobstructed to adopt the upward position. In addition, accessibility of the ACE2 binding site is sterically hindered on the CNTN1-bound RBD^up^, but there is ample space for ACE2 to bind to the remaining RBD’s if they have the upward conformation. Since CNTN1 functions as a co-receptor in combination with ACE2, locking of the adjacent RBD in the down-conformation would reduce availability of spike to ACE2 and inhibit the structural rearrangement needed for membrane fusion. However, opening of the third unobstructed RBD^down^ provides room for the second RBD^down^ to transition upward explaining the avidity effect observed in our interaction experiment. Straightforward modelling of a spike-CNTN1-ACE2 complex, by superposing available structures, indicates that ACE2 and CNTN1 can simultaneously bind a spike trimer that has two or three RBD’s in the upward conformation (Fig. S5).

Newer variants of SARS-CoV-2 spike have emerged since the Wuhan strain used in this study. Considering the most prevalent variants such as the alpha (B1.1.7), beta (B.1.351), gamma (P.1) and omicron (BA.1) variants, mutations at the binding interface are found only in the omicron variant (Fig. 4B). The two mutations in the omicron variant, S373P and S375F, are located next to and on beta strand β2, and introduce a more hydrophobic region (PFF) compared to the Wuhan strain (SFS). These two mutations, together with S371L, stabilize the one-RBD^up^ conformation in the omicron variant, preventing the other RBDs to adopt the upward positions upon ACE2 binding^59^. More generally, compared to the Wuhan strain, subsequent variants show decreased RBD^up^ flexibility^30,60^, which may decrease the accessibility for CNTN1 to bind. While several neuronal infection models are less susceptible to the SARS-CoV-2 omicron variant compared to the D614G and delta variants^61,62^, it remains unclear if CNTN1 plays a role in this difference.

Structures of spike in complex with several antibodies and nanobodies have previously been reported that, based on comparison with the spike-CNTN1 structure, sterically interfere with CNTN1 binding (Fig. 5). Class 4 antibodies have overlapping binding sites with CNTN1 on spike-RBD. Also, class 3 and class 1 antibodies may sterically clash with CNTN1 considering their extended structures. This raises an interesting question about the potential of neutralizing antibodies to mitigate SARS-CoV-2 neurovirulence and the extent to which CNTN1-spike interactions play a role. The effects of monoclonal antibodies on SARS-CoV-2 neurovirulence are, however, not well understood.

In conclusion, we report the affinity of SARS-CoV-2 spike binding to co-receptor CNTN1 and elucidate its general binding interface by presenting a spike cryo-EM map in complex with one CNTN1 molecule. The interface of CNTN1 bound to the RBD^up^ of spike differs from the ACE2 binding site and its placement elicits questions regarding functional rearrangement of spike protomers in combination with CNTN1’s function as a co-receptor of ACE2. These data provide a basis for further targeted research into the relevance of CNTN1 in neuropathogenesis of SARS-CoV-2 and its subsequent variants.

## Data availability

Cryo-EM composite density maps for spike-CNTN1 complex have been deposited to the EMDB with identifier EMD-XXXX, EMD-57277, EMD-57278 and the structural model has been deposited to the PDB with XXXX.

## Acknowledgements

This work benefited from access to the Netherlands Centre for Electron Nanoscopy (NeCEN) at Leiden University, an Instruct-ERIC centre. This research was funded by the Dutch Research Council NWO Gravitation 2013 BOO, Institute for Chemical Immunology (ICI; 024.002.009), as well as EM facility access through NEMI (184.034.014). This work was partially funded by the Corona Accelerated R&D in Europe (CARE) project. The CARE project has received funding from the Innovative Medicines Initiative 2 Joint Undertaking (JU) under grant agreement No 101005077. The JU receives support from the European Union’s Horizon 2020 Research and Innovation Programme, the European Federation of Pharmaceutical Industries and Associations, the Bill & Melinda Gates Foundation, the Global Health Drug Discovery Institute and the University of Dundee. The content of this publication only reflects the author’s views, and the JU is not responsible for any use that may be made of the information it contains.

## Materials & methods

### Expression and purification of CNTN1

IMAGE Clone 30099512 was used as template to obtain constructs for full ectodomain CNTN1 (residues 21-996), CNTN1^Ig1–6^ (residues 21-604), CNTN1^FN1–4^ (residues 604-996) by PCR using primers including a 5’-BamHI restriction site and 3’-NotI restriction site to subclone into HEK293T expression plasmids pUPE107.03 (N-terminal cystatin secretion signal peptide, C-terminal His_6_-tag) or pUPE107.62 (N-terminal cystatin secretion signal peptide, C-terminal biotin acceptor peptide (BAP)-His_6_ tag) provided by ImmunoPrecise Antibodies (IpA) as described earlier^46^. Full ectodomain CNTN1 was expressed by large scale (1L) EBNA1-expressing HEK293-E cells (IpA). After 6 days of expression, cells were spun down by centrifugation at 5000x g at 4 °C for 20 minutes, after which supernatant medium was filtered through a 0.22 µM Steritop vacuum filter (Merck). Fitlered medium was loaded on a 5 mL HisTrap Ni^2+^-excel column (Cytiva) connected to an Äkta system and washed with >20 CV wash buffer (25 mM Hepes pH 7.5, 500 mM NaCl) until UV280 signal remained stable. Subsequently, CNTN1 protein was eluted from the column with elution buffer (25 mM Hepes pH 7.5, 500 mM NaCl, 500 mM Imidazole) and collected in 1 mL fractions. Protein containing fractions were pooled and concentrated using a 50 kDa cut off Amicon Ultra Centrifugal filter (Merck) to a volume of <500 µL. Sample was centrifuged for 10 minutes at 20.000x g and loaded on a Superdex200 Increase SEC column (Cytiva) equilibrated with SEC buffer (25 mM Hepes pH 7.5, 150 mM NaCl). Fractions were loaded on an SDS-PAGE gel to assess purity, and concentration was measured by OD_280_. CNTN1 containing fractions were further concentrated to a final concentration of 3 mg/mL and flash frozen until further use.

### Expression and purification of SARS-CoV-2 6P

A human codon-optimized gene encoding the 6P-stabilized SARS-CoV-2 S ectodomain expression construct^63^ was synthesized by GenScript. The construct comprised spike protein residues 1–1213 from the Wuhan-Hu-1 strain (GenBank: QHD43416.1), followed by a C-terminal T4 foldon trimerization motif, an octa-histidine tag, and a Twin-Strep tag^64^. Spike ectodomains were transiently expressed in HEK-293T cells (American Type Culture Collection (ATCC), CRL-11268) using pCAGGS expression plasmids, and secreted proteins were purified from culture supernatants with streptactin beads (IBA) according to the manufacturer’s instructions. Proteins were eluted in 100 mM Tris-HCl, 150 mM NaCl, 1 mM EDTA, and 2.5 mM D-biotin at pH 8.0.

### SPR measurements

For SPR, CNTN1-BAP-His_6_ constructs were co-expressed with E. coli BirA biotin ligase in pUPE5.02 (N-terminal sub-optimal secretion peptide) in small scale (4 mL) EBNA1-expressing HEK293-E cells with a 20:1 DNA ratio, respectively. A few hours after transfection, 100 µL Tris-HCl buffered sterile biotin (1 mg/mL) was added. After 6 days of expression, cells were spun down at 5000x g for 20 minutes, filtered through a 0.22 µM syringe filter and purified using 100 µL Ni^2+^-excel beads. After bead incubation for 1-2 hours, beads were washed in batch by centrifugating for 5 minutes at 500x g with 60% deceleration and washing with 3× 10 CV wash buffer (25 mM Hepes pH 7.5, 500 mM NaCl). Protein was eluted by adding 100 µL elution buffer (25 mM Hepes pH 7.5, 500 mM NaCl, 500 mM Imidazole), incubating for 15 minutes while rotating and centrifuged to obtain the supernatant to use for chip printing. Samples were diluted with SPR buffer (SEC buffer supplemented with 0.005% Tween-20) to 200, 100 and 50 nM concentrations to print on a P-Strep SensEye® (Ssens) chip using a Continuous Flow Microspotter (CFM, Wasatch Microfluidics). The chip was transferred to a MX96 SPRi instrument (IBIS technologies) and blocked by flowing over 3 mg/mL biotin in SPR buffer. Purified SARS-CoV-2 spike was diluted in SPR buffer and flown over the chip with 4 uL/sec. After every injection, regeneration of the chip was done by injection of 1.5 M MgCl_2_. Data analysis was done using SPRINTX software for local reference subtraction (IBIS Technologies) and Scrubber software, after which a binding curve was fitted to the equilibrium data using a 1:1 Langmuir binding model or a two-state specific binding model for K_D_ calculation.

### Cryo-EM sample preparation, data collection and processing

R1.2/1.3 gold coated on gold 300 mesh grids (QuantiFoil) were glow discharged for 30 seconds at 15 mA before applying 1 mg/mL CNTN1:spike mixture (1:3.15) in a Vitrobot set to humidity of 90% and 4 °C, with a blot time of 4.5 seconds and blot force 0. Grids were rapidly plunge frozen in a liquid ethane/propane mixture. Data was collected with a 300 kV Titan Krios microscope (ThermoFischer Scientific) with a K3 direct electron summit camera (Gatan). Data were collected at 105.000x magnification, resulting in a physical pixel size of 0.836 Å with a total dose of 50 e^−^/Å^2^ over 50 frames. A total of 8226 movies were processed in RELION v5.0.0^65^ (Figure S2). After motion correction with MotionCor2^66^ and CTF estimation with CTFFind^67^, 100 micrographs were used for Autopicking (Laplacian) and extracted particles were 2D classified to generate templates for Topaz^68^ training and picking for the entire dataset. Approximately 2.6 million particles were used for initial 3D classification to select for particles containing one RBD in the up conformation (∼1 million particles). After 3D refinement, particles were subjected to 3D classification into 20 classes without re-alignment using a soft and extended focussed mask on the RBD^up^ domain. Two classes displaying additional CNTN1 density were selected resulting in a total of ∼16.5k particles. In parallel, the initial 1 million particles with one RBD^up^ were subjected to 3D refinement, CTF refinement and polishing to obtain particles with improved signal-to-noise ratio. The ∼16.5k complex particles were reselected from the polished set for subsequent refinement to a nominal resolution of 3.5 Å. The density for the RBD^up^-CNTN1 segment was improved by local refinement to a nominal resolution of 7.2 Å. These two maps were combined into a final volume using ChimeraX^69–71^.

### Model refinement and interpretation

Using ChimeraX, models of spike (PDB ID: 7DWX^72^) and the Ig1-Ig4 domain segment of CNTN1 (PDB ID: 7OL2^73^) were rigid body fitted into the density. Glycans were removed to allow for flexible domain refinement with Namdinator^74^ with the resolution set to 10 Å and 200.000 simulation steps. Interfacing residues were calculated using PDBePISA^75^ web tool and images created with ChimeraX^69–71^.

To determine the displacement of the RBD^up^-CNTN1 combination, two distinct complex classes were 3D refined and the structures were modelled in these maps using Namdinator. Superpose^76^ in the CCP4 package^77^ was used to calculate the displacement of the RBD^up^-CNTN1 combination between the two spike-CNTN1 structures.

