## Supplementary Information for "Structure of SARS-CoV-2 spike in complex with its co-receptor the neuronal cell adhesion protein contactin 1"

- Table S1
- Figures S1-S5

**Table S1.** Data collection and refinement statistics for spike-CNTN1 complex model and cryo-EM composite map.

| **Cryo-EM data collection refinement and validation statistics** | | |
|---|---|---|
| Magnification | | 105kx |
| Voltage (kV) | | 300 |
| Electron exposure ($e^{-}$/ $Å^2$) | | 50 |
| Defocus range (μm) | | -2.5 to -0.8 |
| **Pixel size (Å)** | | 0.836 |
| Symmetry imposed | | C1 |
| Map resolution (Å) | | 3.5 |
| FSC threshold | | 0.143 |
| **Refinement** | | |
| Initial model used | | PDB ID: 7DWX, 7OL2 |
| B-factors ($Å^2$) | | |
| | Protein | 169.71 |
| #non-hydrogen atoms | | 26623 |
| #residues | | 3392 |
| R.m.s. deviations | | |
| | Bond lengths (Å) | 0.008 |
| | Bond angels (degrees) | 1.093 |
| Validation | | |
| | Molprobity score | 2.30 |
| | Clash score | 19.32 |
| | Poor rotamers (%) | 0.7 |
| Ramachandran plot | | |
| | Favored (%) | 91.01 |
| | Allowed (%) | 8.87 |
| | Outliers (%) | 0.12 |

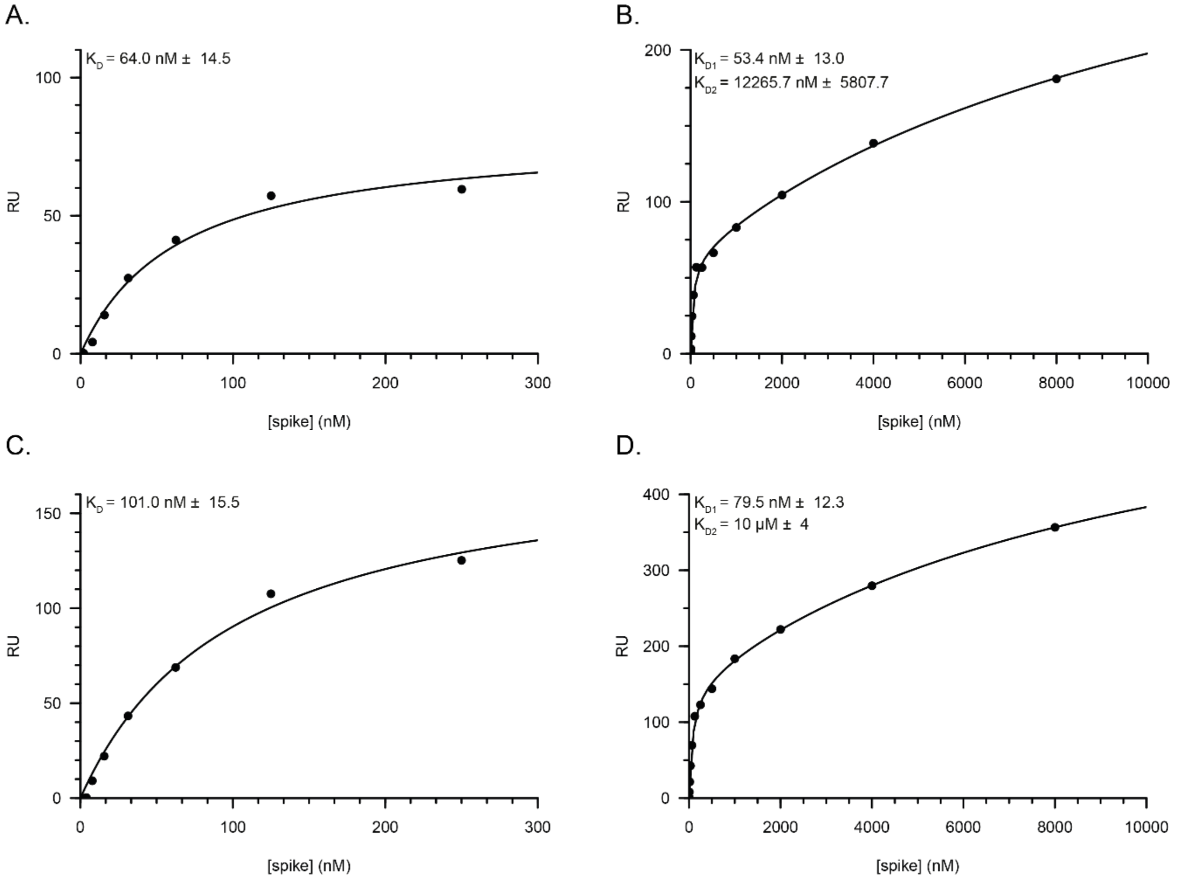


*Figure S1 Replicates of spike-$CNTN1^{fe}$ interaction, as shown in figure 1, show very similar interaction properties and $K_D$ values. A. 1:1 Langmuir binding model fitted to CNTN1 replicate 1, spike concentration ≤ 250 nM. B. Two-state binding model fit of replicate 1, spike concentration ≤ 8 μM. C. 1:1 Langmuir binding model fit CNTN1 replicate 2, spike concentration ≤ 250 nM. D. Two-state binding model fit of replicate 2, spike concentration ≤ 8 μM.*

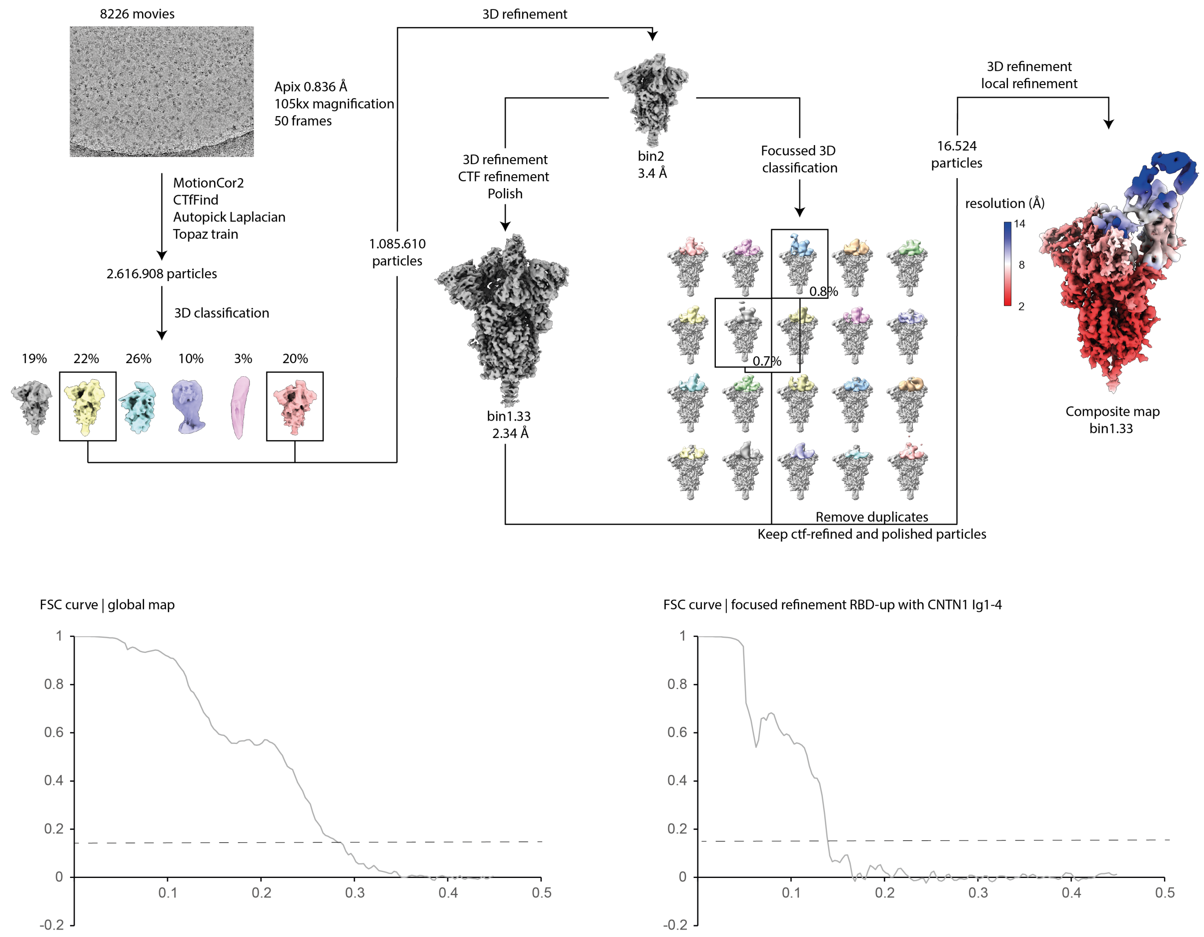


*Figure S2 CryoEM RELION processing scheme of SARS-Cov-2 spike in complex with CNTN1. Dashed lines in FSC curves are at 0.143.*

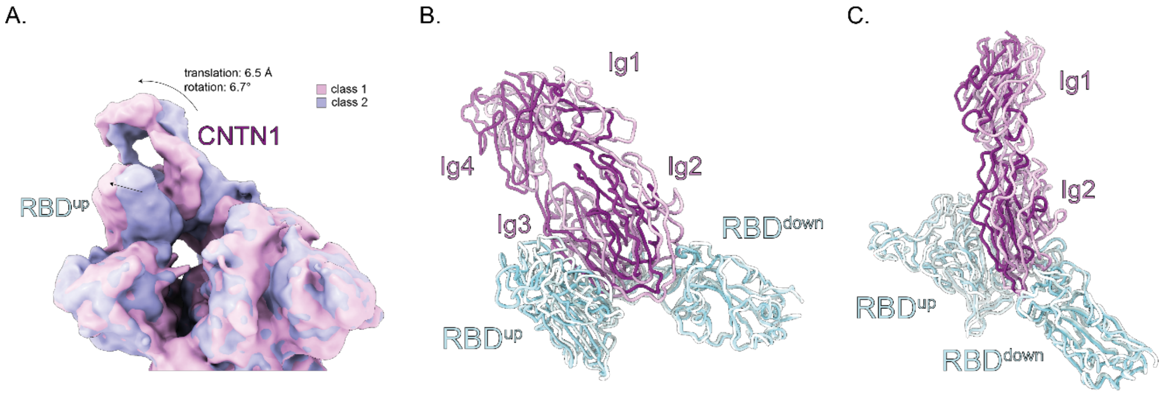


*Figure S3 Dynamic flexibility of CNTN1:RBD$^{up}$ with respect to the rest of spike. A. 2 Gaussian filtered density maps obtained after focused 3D classification. Movement, a 6.5 Å translation and 6.7° rotation, of the CNTN1:RBD$^{up}$ combination is indicated by arrows. B. Models refined against density maps mentioned in A. obtained with Namdinator[74], with equivalent domains represented in similar lighter colours.*

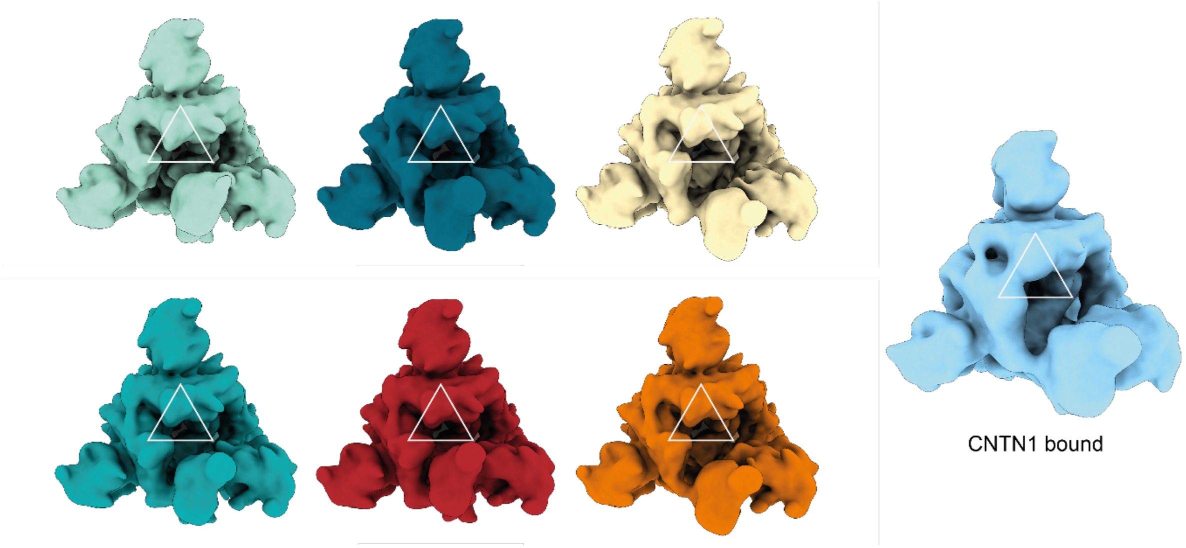


*Figure S4 The RBD$^{up}$ domain has flexibility with respect to the rest of the spike trimer. Gaussian filtered density maps obtained from focussed 3D classification of all spike particles with one RBD$^{up}$. A gaussian filtered volume of the complex density with removed CNTN1 density is displayed in light blue for comparison and shows the confromation of RBD$^{up}$ fits best with that of conformation 6 displayed in orange.*

A.

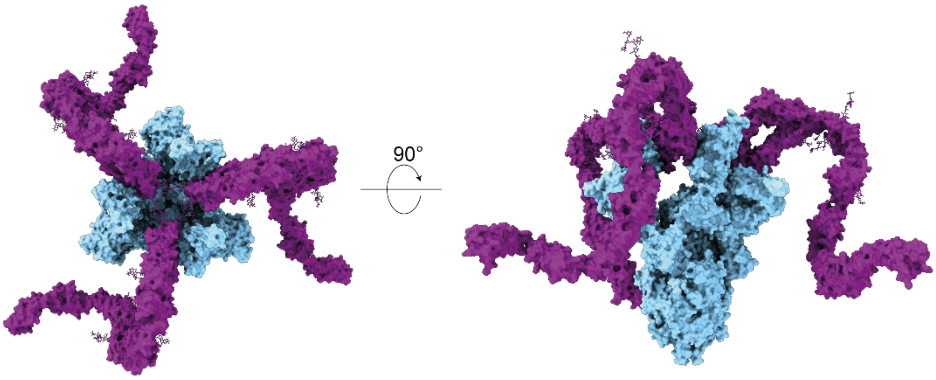


B.

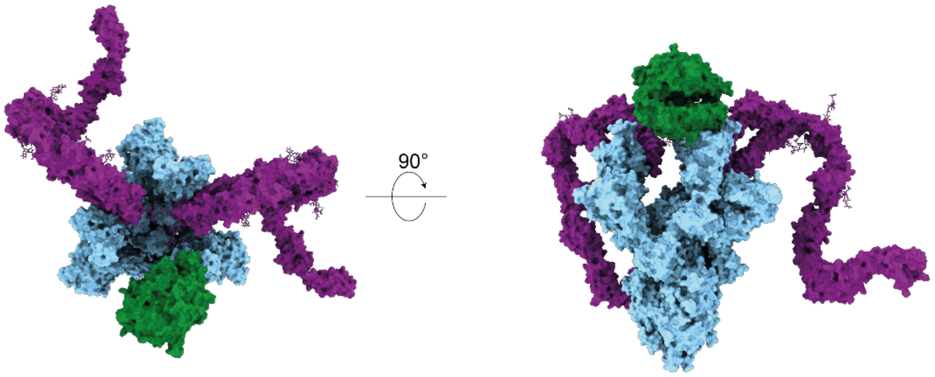


*Figure S5 Models representing multivalent binding of three CNTN1 molecules and mixed co-receptor binding CNTN1 and ACE2 to SARS-Cov-2. PDB ID used for spike: 7R40[48], ACE2: 6VW1[32] and a SAXS based full-ectodomain model of CNTN1[46] A. Model obtained by superposition of three CNTN1 molecules (magenta) to spike (light blue) with three $RBD^{up}$. Note that CNTN1 has flexibility in the Ig4-Ig5 connection and Ig5-FNIII tail that is not modelled. B. Model obtained by superposition of two CNTN1 molecules (magenta) and one ACE2 molecule (green) to spike (light blue). In none of the models steric hindrance is apparent suggesting that these interactions are possible.*
